# A system for CBAF reconstitution reveals roles for BAF47 domains and BCL7 in nucleosome ejection

**DOI:** 10.1101/2021.10.26.465931

**Authors:** Timothy S. Mulvihill, Mary L. Nelson, Naveen Verma, Kevin B. Jones, Bradley R. Cairns

## Abstract

Canonical BAF (CBAF) is an essential 12-protein chromatin-remodeling complex that slides and/or ejects nucleosomes using the alternative catalytic ATP-dependent DNA translocases BRG1 or BRM. Currently, the regulation of BRG1/BRM activity and nucleosome ejection remain incompletely understood. To address this, we developed a system for full CBAF reconstitution and purification, and created a novel nucleosome ejection assay. ARID1A and DPF2 were dispensable for assembly and chromatin remodeling activity, contrasting with prior work. The actin-related protein BAF53A and β-actin components interacted and enhanced DNA translocation, and were required for BCL7A incorporation, which potentiated ejection. BAF47 also regulated ejection, utilizing two stimulatory domains and an autoinhibitory domain. Finally, we provide evidence for ‘direct’ nucleosome ejection at low nucleosome density on closed circular arrays. Taken together, we provide powerful new tools for CBAF mechanistic investigation and reveal new roles for several CBAF components.

## INTRODUCTION

BAF family chromatin remodelers (also termed human SWI/SNF family) are large, multi-subunit complexes that modify chromatin structure to provide DNA access to transcription factors^1,2^. BAF remodels chromatin structure by using the energy of ATP hydrolysis to mobilize nucleosomes—the fundamental repeating unit of chromatin—through two primary mechanisms: linear mobilization of nucleosomes along the DNA, termed sliding, and disassembly of nucleosomes, termed ejection^1,3^. The human BAF family of chromatin remodelers is divided into three subfamilies, Canonical BAF (CBAF), Polybromo-associated BAF (PBAF), and GLTSCR1-associated BAF (GBAF, also known as ncBAF), which share a set of core proteins but are defined by the inclusion of subfamily-specific subunits^4^.

BAF subunits are mutated in nearly 20% of all cancers as well as a number of neurodevelopmental disorders^5–7^. While previous work has characterized a small number of these mutations, the effects of the vast majority of disease-associated BAF mutations on the targeting and/or enzymatic activity of the complex remain unclear, due to our lack of knowledge regarding the roles of certain specific subunits in regulating BAF chromatin remodeling^8–11^. The main barriers to the investigation of BAF regulation are its size and complexity. CBAF, for example, is over 1 MDa in size and comprised of 12 proteins, which (including paralog alternatives) are encoded by a total of 22 genes, with 1,296 possible combinations of subunits^4^.

Previous investigations into the regulation of BAF enzymatic activity have relied on purification of endogenous complexes^12–14^ or reconstitution of partial complexes^15,16^, and been highly informative. However, due to heterogeneity, results obtained by these approaches reflect an ensemble measurement of complexes with a variety of compositions, making it difficult to isolate the effects of a single alteration. Additionally, reconstitution of partial complexes may result in the inadvertent exclusion of important regulatory subunits, potentially skewing results.

Enzymatic activity of CBAF is provided by one of two mutually exclusive ATP-dependent DNA translocases, BRG1 or BRM. Insight into the mechanism of chromatin remodeling has come from studies of related remodelers and homologues from different species, including yeast, flies, and mice^17^. A key unifying feature of remodelers is that ATP hydrolysis is linked to DNA translocation, which involves the processive inchworming of two RecA-like ATPase lobes along the DNA sugar-phosphate backbone, at a rate of one nucleotide per ATP^18,19^, in a manner similar to bacterial DNA translocases for DNA repair^20^. For BAF-related remodelers, DNA translocation from within the nucleosome—resulting from the RecA-like lobes residing two superhelical turns from the dyad—while other domains anchor on the histone octamer, enables both nucleosome sliding and ejection^18,21,22^. Sensitive and scalable assays to assess each of these activities are essential for understanding how different subunits regulate CBAF activity on a mechanistic level.

Here, advancing on prior work with partial BAF sub-complexes^15,16,23^, we develop a system for the production and purification of recombinant full CBAF complex from human cells. We use the system to investigate assembly dependencies of specific subunits, the roles of particular subunits in regulating enzymatic activity, and the fundamental mechanism of nucleosome ejection—through the additional development of a new assay for assessing ejection.

## RESULTS

### Expression and purification of recombinant CBAF from human cells

Our goal was a versatile and efficient system to produce and isolate full 12-protein CBAF, or any CBAF mutant derivative, for biochemical or structural studies. We advanced on prior work on partial BAF complexes^15,16,23^ by adapting an existing cloning system, biGBac, for use in mammalian cells by replacing the entry vector, pLib, with pLibMam, a pFastBac1 derivative with a strong constitutive CMV promoter replacing the polh promoter (Fig. 1a)^24^. The predefined oligonucleotides used in the biGBac system were redesigned to make the pLibMam vector compatible with the biGBac shuttle vectors by changing the priming portion of each oligonucleotide to match the pLibMam vector, while retaining the Gibson homology sequences that match the biGBac pBig1 vectors. This same logic could in principle be used to make any expression vector compatible with the biGBac system.

**Fig. 1.**
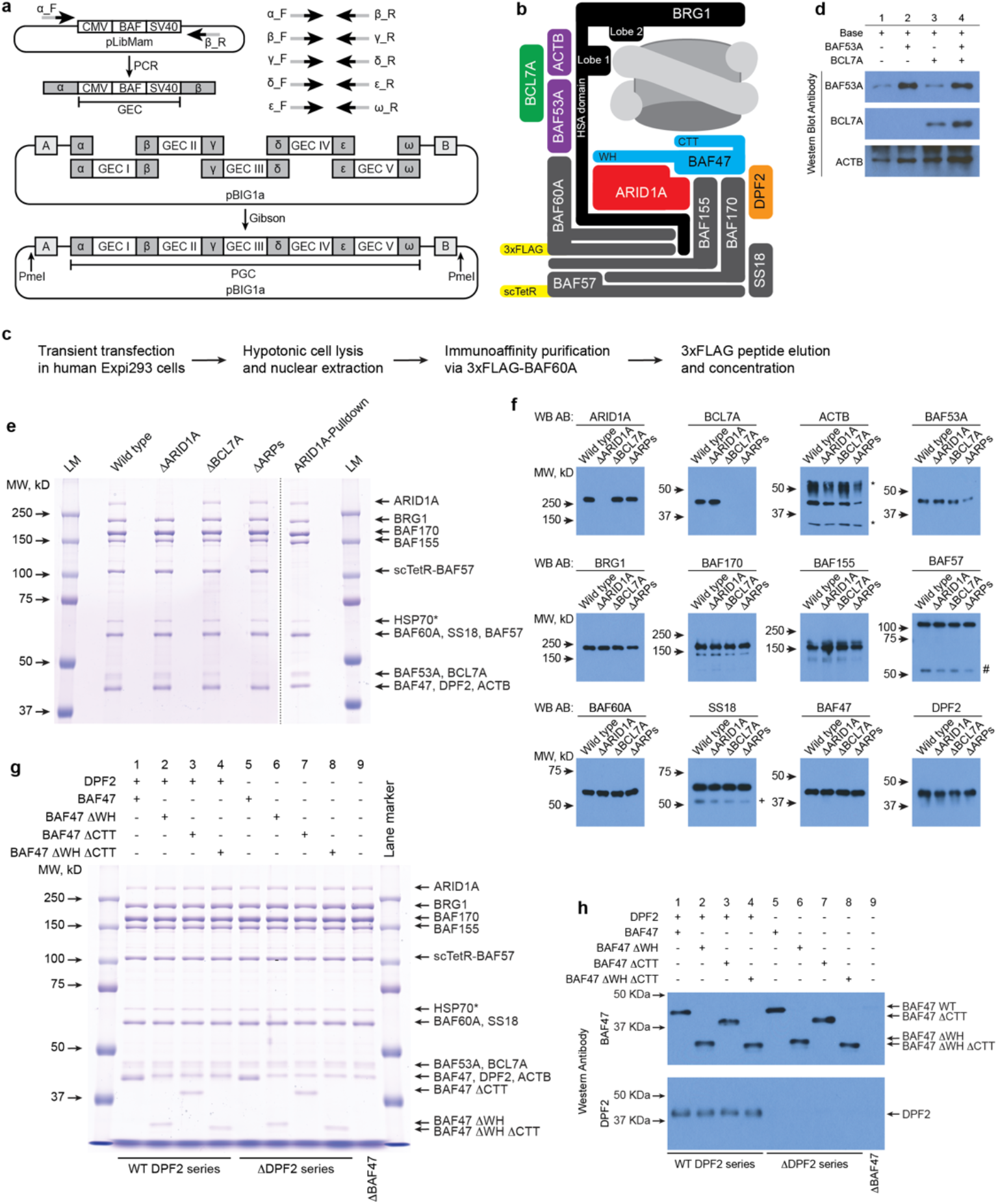
A system for the production and purification of fully-assembled recombinant CBAF complexes from human cells. **a**, Schematic of the modified biGBac cloning system. CBAF subunits (BAF) are cloned into the pLibMam vector. Predefined oligonucleotides are used to PCR amplify a gene expression cassette (GEC) while introducing Gibson homology sequences (a, β etc.). Gibson assembly is used to combine GECs with a biGBac pBIG1 vector. Digestion of pBIG1 vectors with PmeI releases a poly-gene expression cassette (PGC) flanked by additional Gibson homology sequences (A, B) to subclone into a pBIG2 vector. **b**, Schematic of the recombinant wild type CBAF complex produced in this study with key features highlighted. **c**, Protocol for the expression and purification of recombinant CBAF from human cells. **d**, Western blots for ARP module assembly dependencies. Base expression vectors included in all transfections were pBIG2-ΔARPs and pLibMam-ARID1A. pLibMam-BAF53A, pLibMam-BCL7A, neither, or both were co-transfected as indicated. **e**, Coomassie-stained SDS-PAGE gels of purified recombinant CBAF complexes. See Methods for details of plasmid transfections. HSP70* is a low-level contaminating protein that is not part of CBAF, but is common in large-scale expression/purification systems, as it binds to various partially-folded regions. LM: Lane Marker. **f**, Western blots of purified recombinant CBAF complexes shown in panel **e**. *: IgG heavy and light chain. #: endogenous BAF57. +: endogenous short splice variant of SS18. **g**, Coomassie-stained SDS-PAGE gels of purified recombinant CBAF complexes. See panel **e** for HSP70* explanation. **h**, Western blots of purified recombinant CBAF complexes shown in panel **g**.

We then assembled a wild type (WT) CBAF expression vector containing the genes encoding BRG1, BAF170, BAF155, BAF57, BAF60A, SS18, BAF53A, BCL7A, BAF47, and DPF2 (Fig. 1b). Due to our interest in examining roles for ARID1A, ARID1A was expressed from a separate vector (pLibMam-ARID1A). Finally, β-actin was not overexpressed, as endogenous sources sufficed. Here, the most widely expressed paralog of each subunit was selected for inclusion in the system (see Methods). To aid in purification of recombinant CBAF, a 3xFLAG tag was added at the N-terminus of BAF60A. A single-chain version of the DNA binding domain of the tetracycline repressor (scTetR) was added at the N-terminus of BAF57 to allow tethering of complexes to the TetO DNA sequence to enable a Tet-tethered DNA translocation assay, and to eliminate the need for a double affinity purification to select for TetR heterodimers^25^. The expression vectors were transiently transfected into human Expi293F cells, and recombinant CBAF was purified from nuclear extracts via 3xFLAG immunoaffinity purification, competitively eluted with 3xFLAG peptide, concentrated, aliquoted, and frozen (Fig. 1c).

We first investigated the roles of the Actin-Related Protein (ARP) module in regulating CBAF enzymatic activity. Previous studies identified the ARP module of RSC, a yeast homolog of BAF, as a key regulatory module^26^ that enhances the efficiency with which ATP hydrolysis is ‘coupled’ to DNA translocation, increasing sliding efficiency and potentiating nucleosome ejection^27^. However, whether this regulatory function is evolutionarily conserved in CBAF is not known, as CBAF contains the actin-related protein BAF53 and β-actin itself, rather than two ARPs as in RSC^28,29^. As expected, given that β-actin and BAF53 form an obligate heterodimer within CBAF, exclusion of BAF53A from the expression vector led to a substantial reduction in both subunits in the resulting purified complex. Unexpectedly, exclusion of BAF53A also greatly reduced BCL7A incorporation, indicating that BCL7A is a member of the CBAF ARP module, in keeping with its physical location within CBAF (Fig. 1d)^30^. Therefore, to thoroughly test the role of the ARP module in regulating CBAF enzymatic activity (below) we purified both a complex lacking only BCL7A (ΔBCL7A) and a complex lacking BCL7A, BAF53A, and β-actin (ΔARP) (Fig. 1e, f). Purified complexes were assessed with analytical size exclusion chromatography and found to be 85-89% homogenous and monodisperse (Supplementary Fig. 1).

As a second test of the system, we investigated the role of ARID1A in CBAF assembly. Currently, there are conflicting reports regarding the requirement of an ARID1 paralog for CBAF stability and activity. Multiple reports indicate its requirement for stability^4,31^ while others claim dispensability^15,32^. Similarly, ARID1A loss has been reported to cause a significant decline in nucleosome remodeling activity^15^, yet reconstituted partial BAF complexes lacking ARID1A are active^16^. Notably, we found that CBAF readily assembled in the absence of ARID1A (Fig. 1e, f). To confirm that ARID1A is present in stoichiometric levels in our WT CBAF complex, we purified a version of the complex with a His-tag on ARID1A, allowing us to perform a second pulldown of ARID1A-containing complexes after the 3xFLAG peptide elution step. The relative abundance of each subunit in this complex closely matched the WT purification, confirming that ARID1A is present at stoichiometric levels in the WT complex, but simply stains poorly with Coomassie Blue (Fig. 1e).

As a third test of the system, we examined the role of BAF47 (and domains within) along with DPF2 in regulating CBAF enzymatic activity. BAF47 is frequently mutated in cancer, most notably malignant rhabdoid tumors, which are characterized by biallelic inactivation of *SMARCB1*, the gene encoding BAF47^33,34^. The C-terminal tail (CTT) of Sfh1, the yeast homolog of BAF47, has been identified as a key domain that contacts the nucleosome acidic patch and potentiates nucleosome ejection^35^. Mutations in the BAF47 CTT are associated with various developmental disorders and have been shown to compromise CBAF chromatin remodeling activity^13^. Mutations in the N-terminal winged helix (WH) domain of BAF47 are associated with Schwannomatosis^36^, but there are conflicting reports regarding its location within the complex^15,30^. BAF47 also contains two centrally located RPT domains that are required for its association with the complex as well as for incorporation of DPF2^30^. To investigate the roles of each of these domains we produced a set of nine complexes to test the effects of loss of the BAF47 WH, BAF47 CTT, and DPF2 alone and in combination, as well as deletion of both subunits entirely (Fig. 1g, h).

### A quantifiable assay for nucleosome ejection using closed circular arrays

As nucleosome ejection is linked to ARP and BAF47 function, we next developed a new, scalable, and quantifiable method of assessing nucleosome ejection activity on defined DNA templates based on existing assays^12,27,37^. The assay is predicated on three key principles: 1) plasmid DNA purified from bacteria is inherently negatively supercoiled, 2) nucleosomes store negative supercoils (1 per nucleosome) and protect them from relaxation by *E. coli* topoisomerase I (Topo I), and 3) plasmid topoisomers can be readily distinguished on an agarose gel. With these principles in mind, we assembled poly-nucleosome arrays on a plasmid containing 12 repeats of a 200bp Widom 601 nucleosome positioning sequence, and then incubated the assembled arrays with CBAF and Topo I. As CBAF ejects nucleosomes, negative supercoils are released into the plasmid and relaxed by Topo I, leading to a reduction in linking number and decreased electrophoretic mobility on an agarose gel (Fig 2a, b). We note that this assay lacks histone chaperones or free DNA acceptors of histones, which can assist the ejection process, but were omitted to focus on the mechanics of CBAF ejection in isolation.

**Fig. 2.**
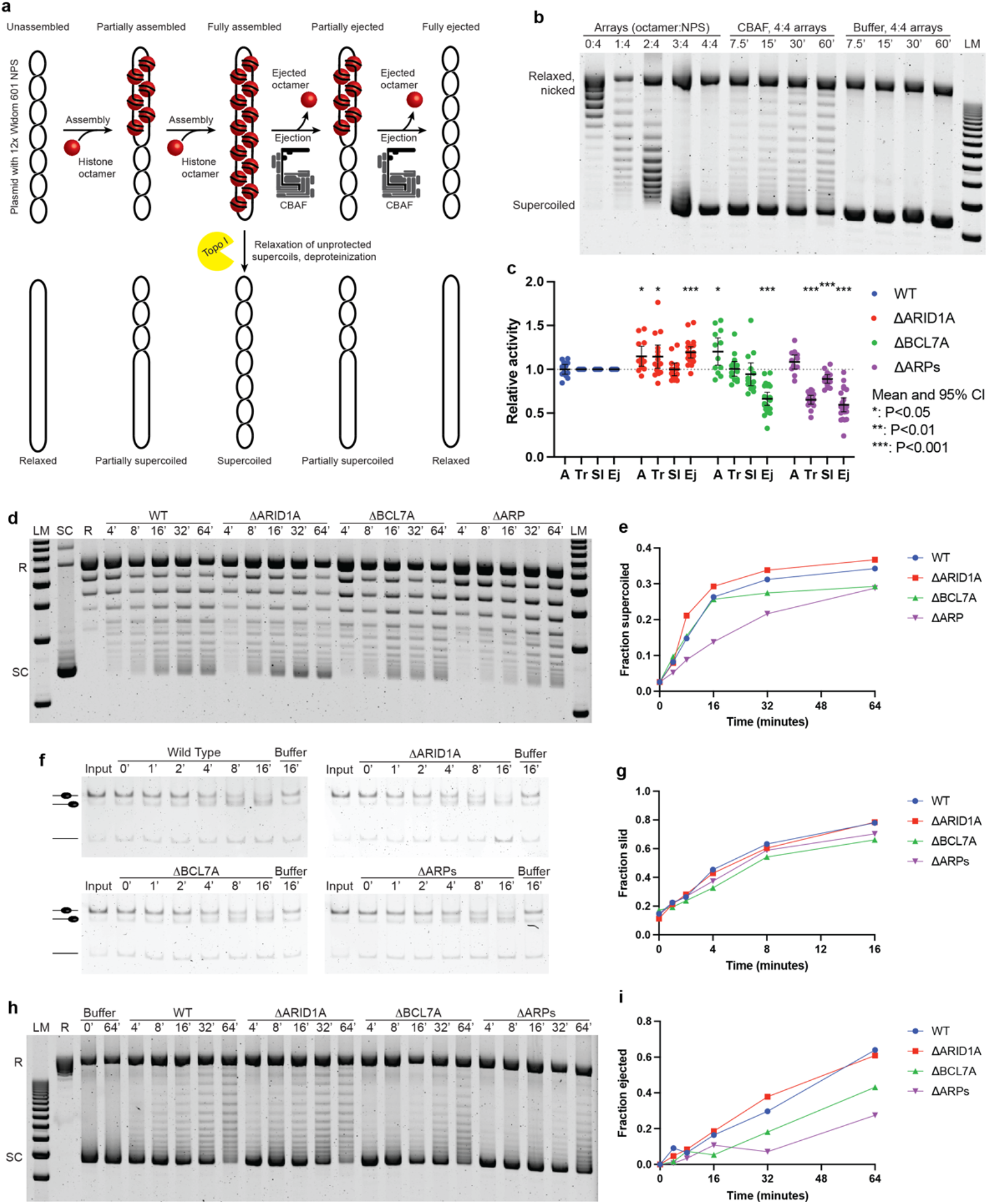
ARPs and BCL7A enhance chromatin remodeling through distinct mechanisms, while ARID1A is dispensable for remodeling. **a**, Schematic of the nucleosome ejection assay. **b**, Example of an ejection assay gel. Arrays were assembled with various molar ratios of octamer to nucleosome positioning sequence (NPS) prior to treatment with Topo I. 4:4 arrays were incubated with CBAF or buffer for the indicated times. LM: Lane Marker. **c**, Summary of data from ATPase (A), translocation (Tr), sliding (Sl) and ejection (Ej) assays. Error bars represent mean +/- 95% confidence interval. **d**, Representative gel for a DNA translocation assay. Tet-tethered DNA translocation on a plasmid containing the TetO sequence in the presence of Topo 1 results in the accumulation of positive supercoils as translocation occurs. R: Relaxed plasmid; SC: Supercoiled plasmid; LM: Lane Marker. **e**, Quantification of the DNA translocation assay shown in panel **d**. The fraction of total lane intensity representing supercoiled topoisomers is quantified and plotted over time. **f**, Representative gel for a nucleosome sliding assay. **g**, Quantification of the nucleosome sliding assay shown in panel **f**. The fraction of end-positioned mononucleosomes (slid) are quantified and plotted over time. **h**, Representative gel for a nucleosome ejection assay. R: Relaxed plasmid; SC: Supercoiled plasmid; LM: Lane Marker. **i**, Quantification of the nucleosome ejection assay shown in panel **h**. The fraction of total lane intensity corresponding to fully supercoiled topoisomers is quantified and normalized to the Buffer T=0’ timepoint. The resulting values are subtracted from 1 and plotted over time.

### ARPs and BCL7A, but not ARID1A, enhance chromatin remodeling

We used this nucleosome ejection assay, along with ATPase, DNA translocation, and nucleosome sliding assays to investigate the roles of ARID1A and the ARP module in regulating CBAF enzymatic activity. DNA-dependent ATPase activity is assayed using a colorimetric assay that measures the inorganic phosphate released by ATP hydrolysis by complexation with molybdate-malachite green. DNA translocation is assessed with a topology assay that measures supercoiling induced by translocation along the DNA sugar-phosphate backbone while tethered to a fixed location on the plasmid via the TetR-TetO interaction. Nucleosome sliding is assayed by the repositioning of a centrally located nucleosome to the end of a 200bp Widom 601 nucleosome positioning sequence. Each assay was performed four or more times—at least twice with each of two different purified complexes for each variant—with at least three timepoints or replicate samples in each experiment used for statistical analysis. A summary of the data, normalized to WT activity levels, is shown (Fig. 2c).

First, DNA-dependent ATPase activity (at Vmax) was not decreased by loss of ARID1A, BCL7A, or the ARP module. This demonstrates that the core ATPase activity of BRG1 remains intact in each of these complexes and that reductions in activity observed in the other assays are due to defects in linking ATPase activity to particular remodeling outcomes, rather than a defect in the motor itself. While the ΔBCL7A complex had WT levels of DNA translocation, the ΔARPs complex displayed significantly reduced activity (Fig. 2d, e, and Supplementary Fig. 2), consistent with previous work on RSC and supporting an evolutionarily conserved role for ARPs in improving the efficiency with which ATP hydrolysis is coupled to DNA translocation^27^. As expected, given their relative DNA translocation capacities, the ΔBCL7A complex had WT levels of nucleosome sliding while the ΔARPs complex had reduced nucleosome sliding activity (Fig. 2f, g, and Supplementary Fig. 3). Intriguingly, the defect in nucleosome sliding observed with loss of the ARP module was smaller in magnitude than the defect in DNA translocation. This may reflect differences in the biophysical parameters of the two assays. The tethered DNA translocation assay requires high processivity (10bp per supercoil) and considerable force resistance in its ATPase-DNA ‘grip’ to produce highly supercoiled topoisomers. In contrast, nucleosome sliding is achievable with low processivity and low force resistance due to the requirement of breaking only 1-2 histone-DNA contacts at a time^1^.

Interestingly, despite approximately WT levels of ATPase, translocation, and sliding activity, the ΔBCL7A complex had a significant reduction in nucleosome ejection (Fig. 2h, i, and Supplementary Fig. 4). Nucleosome ejection is a high-force, high-processivity action, as it requires simultaneous rupture of multiple histone-DNA contacts. Because the robust translocation activity of this remodeler indicates that processivity and DNA grip is intact, we speculate that loss of BCL7A causes a reduction in force resistance due to a defect in anchoring on the histone octamer and/or reduced DNA translocation processivity on the octamer. As expected, given its reduced DNA translocation and nucleosome sliding activities, the ΔARPs complex also had significantly decreased nucleosome ejection activity. While a portion of the loss of ejection activity can be attributed to loss of BCL7A, the ΔARPs complex trended towards lower levels of nucleosome ejection than the ΔBCL7A complex. Here, the trend towards decreased activity relative to the ΔBCL7A complex suggested that both the presumed histone anchoring provided by BCL7A and the enhanced coupling provided by BAF53A and β-actin may be required for full WT levels of nucleosome ejection.

Surprisingly, the ΔARID1A complex was found to have WT or increased levels of ATPase, DNA translocation, nucleosome sliding, and nucleosome ejection (Fig. 2c-i). This result contrasts with a previous report that showed a major decrease in nucleosome sliding activity with loss of ARID1A^15^. While both this study and the previous report used the Widom 601 nucleosome positioning sequence as a template for sliding assays, the previous result was obtained using partial CBAF complexes lacking both SS18 and BCL7A—a subunit shown above to play a key role in regulating nucleosome remodeling. The prior study also used *Xenopus* histone octamers, rather than the *Drosophila* octamers used in this study. Histone octamer identity has been shown to influence chromatin remodeling by RSC, raising the possibility that ARID1A loss reduces remodeling activity only in particular nucleosome contexts^38^.

### BAF47 regulates ejection through inhibitory and stimulatory domains

We next investigated the roles of BAF47 and DPF2. Here, each assay was performed four times, generally twice with each of two different purified complexes for each variant, with multiple timepoints or replicates from each experiment used for statistical analysis, as summarized (Fig. 3a). The nine complexes fall into two series (+DPF2 and ΔDPF2), with four different BAF47 variants (WT, ΔWH, ΔCTT, and ΔWH ΔCTT) in each, along with a ninth complex (ΔBAF47 ΔDPF2). All variant complexes trended toward decreased ATPase activity, with four (BAF47 ΔCTT +DPF2, BAF47 WT ΔDPF2, BAF47 ΔWH ΔCTT ΔDPF2, and ΔBAF47 ΔDPF2) reaching statistical significance. However, all had at least 75% of WT activity, indicating that the core ATPase activity of BRG1 is only modestly affected by each of these alterations.

**Fig. 3.**
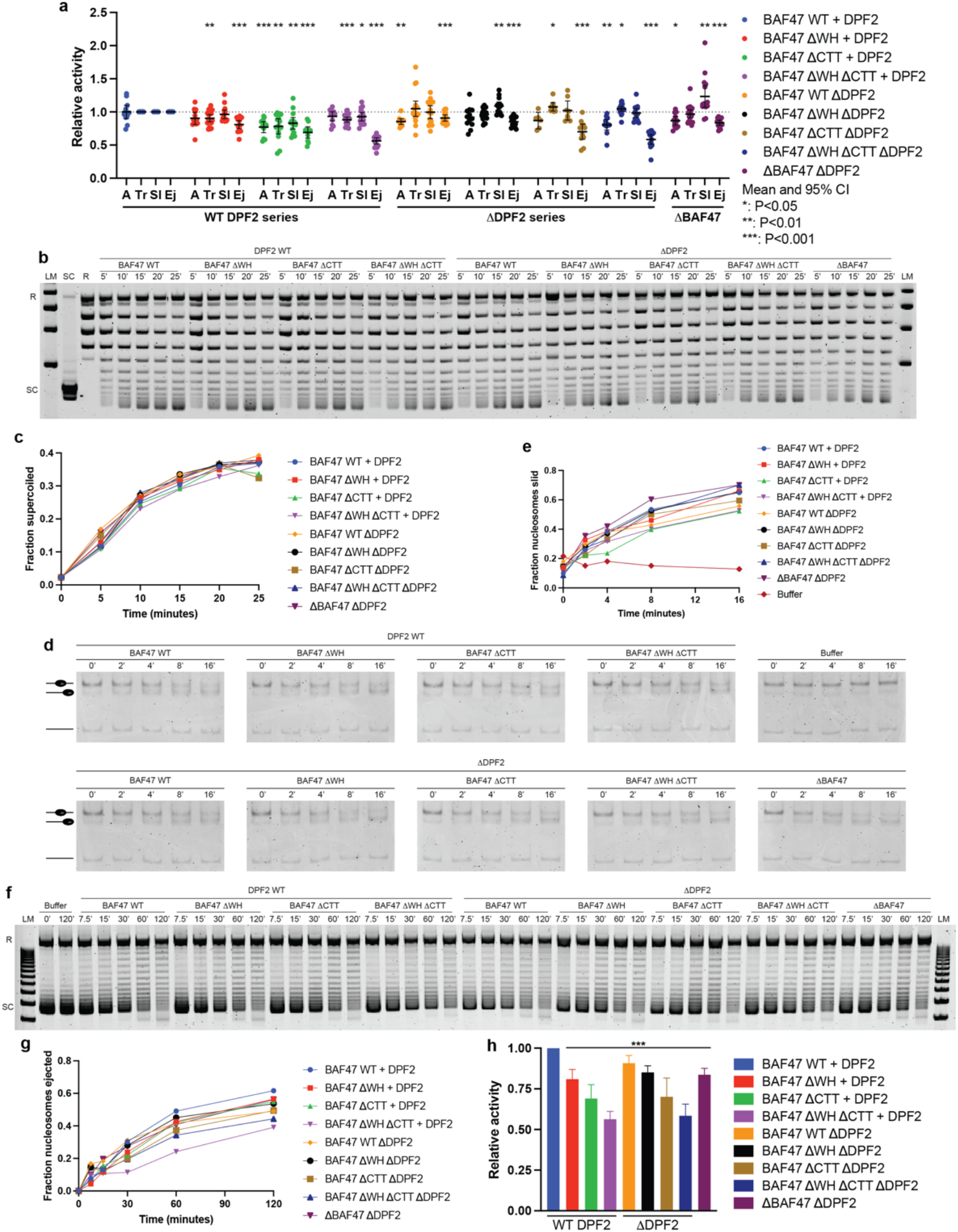
BAF47 regulates nucleosome ejection through multiple regulatory domains. **a**, Summary data from ATPase (A), translocation (Tr), sliding (Sl) and ejection (Ej) assays as in Fig. 2. Error bars represent mean +/- 95% confidence interval. **b**, Representative gel for a DNA translocation assay as in Fig. 2. R: Relaxed plasmid; SC: Supercoiled plasmid; LM: Lane Marker. **c**, Quantification of the DNA translocation assay shown in panel **b**. The fraction of total lane intensity representing supercoiled topoisomers is quantified and plotted over time. **d**, Representative gel for a nucleosome sliding assay. **e**, Quantification of the nucleosome sliding assay shown in panel **d**. The fraction of end-positioned mononucleosomes (slid) are quantified and plotted over time. **f**, Representative gel for a nucleosome ejection assay. R: Relaxed plasmid; SC: Supercoiled plasmid; LM: Lane Marker. **g**, Quantification of the nucleosome ejection assay shown in panel **f**. The fraction of total lane intensity corresponding to fully supercoiled topoisomers is quantified and normalized to the Buffer T=0’ timepoint. The resulting values are subtracted from 1 and plotted over time. **h**, Relative ejection activity of CBAF variants with BAF47 truncations and/or DPF2 deletion. Error bars represent mean +/- 95% confidence interval. Significance was calculated using a paired t-test of each variant complex relative to WT. ***: P≤0.001.

All variant complexes in the +DPF2 series had significantly reduced DNA translocation activity (Fig. 3b-c, and Supplementary Fig. 5). However, the observed decreases closely matched the respective reductions in ATPase activity, indicating that the translocation deficits can likely be attributed to reduced ATPase activity rather than a reduction in coupling efficiency. The variant complexes in the ΔDPF2 series all had DNA translocation capacities that exceeded their relative ATPase activities, indicating that coupling efficiency is intact or slightly elevated in each of the variant complexes. In addition, the ΔBAF47 ΔDPF2 complex had WT levels of DNA translocation. Collectively, these results indicate that neither DPF2 nor BAF47 loss confers a deficit in processivity or force resistance.

Nucleosome sliding activity was highly correlated with DNA translocation activity for all variant complexes in the +DPF2 and ΔDPF2 series (Fig. 3d, e, and Supplementary Fig. 6). Two complexes (BAF47 ΔCTT +DPF2 and BAF47 ΔWH ΔCTT +DPF2) had statistically significantly reduced nucleosome sliding activity, although as with the DNA translocation assay, the observed reduction can be attributed to decreased ATPase activity rather than a specific defect in nucleosome sliding. Intriguingly, the ΔBAF47 ΔDPF2 complex displayed significantly increased nucleosome sliding activity. Taken together, these results indicate that neither DPF2 nor BAF47 loss confers a deficit in nucleosome engagement.

Notably, all variant complexes had a statistically significant decrease in ejection activity, with the magnitude of the decrease being attributable to the specific BAF47 truncation, which was consistent across both the +DPF2 and ΔDPF2 series (Fig. 3f-h, and Supplementary Fig. 7). Specifically, deletion of the BAF47 WH caused a ~15-20% reduction in activity, deletion of the CTT caused a ~30% reduction in activity, and deletion of both the WH and the CTT had an additive effect, causing a ~40-45% reduction in activity. Importantly, for all BAF47 truncations, the measured decrease in ejection activity exceeded the decrease in activity in each of the other assays, indicating a specific role for BAF47 in regulating nucleosome ejection, rather than remodeling in general. The ejection activity of each BAF47 variant was unaffected by deletion of DPF2, indicating that DPF2 does not regulate nucleosome ejection. Surprisingly, deletion of both BAF47 and DPF2 conferred a ~15-20% reduction, less severe than the combined WH and CTT truncations, implying the presence of a previously unknown autoinhibitory domain in the central portion of BAF47.

### CBAF is capable of direct ejection of nucleosome

Finally, we adapted our new nucleosome ejection assay to garner insight into the mechanism of nucleosome ejection by CBAF. Two possible mechanisms have been proposed—direct ejection of nucleosomes^39^ and unspooling of DNA from an adjacent nucleosome^40–42^. Direct nucleosome ejection involves ejection of the nucleosome that is directly engaged/bound by the remodeler, whereas the spooling mechanism involves first sliding a nucleosome into an adjacent nucleosome, followed by additional DNA translocation/sliding, resulting in unspooling of DNA from the neighboring octamer—causing ejection. Importantly, the direct ejection method would allow for ejection of all nucleosomes from a circular array, whereas the spooling mechanism would result in retention of a single nucleosome on the array. We note that these mechanisms are not mutually exclusive, and may instead be regulated and/or occur at very different rates.

Our assessment recognized that the two terminal products of direct ejection or spooling would result in different plasmid topoisomers (with linking numbers of 0 and −1, respectively) suggesting that they might be distinguishable on an agarose gel. However, the presence of nicked DNA in our samples, which migrates at a very similar rate to the fully relaxed plasmid, prevented visualization of the relaxed plasmid and made it challenging to observe whether the final end product of the ejection assay was a fully relaxed plasmid or a topoisomer with a linking number equal to −1. To overcome this limitation, we utilized T5 exonuclease, which degrades nicked—but not covalently closed—plasmids to remove nicked DNA from the assay, allowing for visualization of changes in the relative abundance of the fully relaxed and −1 topoisomers during a timecourse ejection assay (Fig. 4a, b). With this modified ejection assay we observed a clear increase in fully-relaxed plasmid abundance over time, with no observable buildup of a −1 intermediate—thus providing evidence for direct ejection of nucleosomes by CBAF. This result strongly supports a direct ejection mechanism at low nucleosome densities, while leaving open the possibility of a spooling mechanism at high nucleosome densities.

**Fig. 4.**
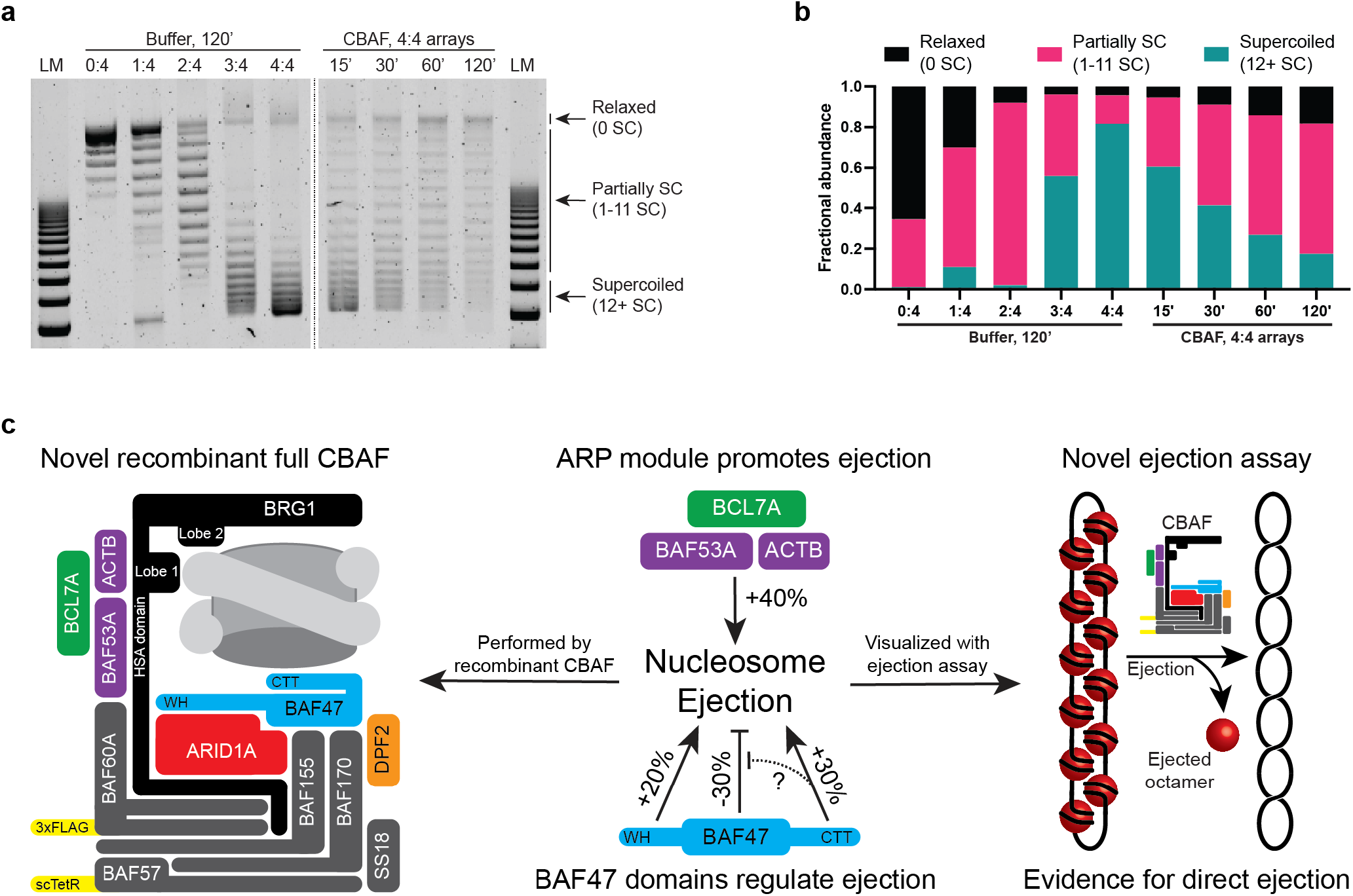
Evidence for direct ejection of nucleosomes from nucleosome arrays by CBAF. **a**, Evidence for direct nucleosome ejection by CBAF. Polynucleosome arrays were assembled with various histone octamer to Widom 601 nucleosome positioning sequence molar ratios (0:4, 1:4, 2:4, 3:4, or 4:4) as indicated. Assembled arrays were incubated with buffer and Topo I for 120’. 4:4 arrays (fully assembled) were used as input for a timecourse ejection assay. All samples were treated with T5 exonuclease to remove nicked plasmid immediately prior to agarose gel electrophoresis. SC: Supercoiled; LM: Lane Marker. **b**, Quantification of the ejection assay shown in panel **a**. The fractions of total lane intensity corresponding to fully relaxed plasmid (0 SC), partially relaxed/supercoiled plasmid (1-11 SC), and fully supercoiled plasmid (12+ SC) are graphed. **c**, A model of the system and results. Purified recombinant CBAF nucleosome ejection activity measured with a novel assay reveals roles for the Actin-Related Protein (ARP) module, including BCL7A, as well as the Winged Helix (WH) and C-Terminal Tail (CTT) domains of BAF47 in regulating ejection activity. See main text for discussion.

## DISCUSSION

Here, we have created a flexible system for the production, purification, and assessment of full recombinant CBAF to facilitate investigation of CBAF assembly, activity, and regulation (Fig. 1a, b). The system allows for the rapid production of WT and variant complexes with defined composition from human cells, to promote physiologically-relevant folding and post-translational modification of subunits. Notably, incorporation of endogenous subunits was minimal (β-actin excepted), owing to the high level of overexpression achieved with the system. This makes the system suitable for production and isolation of variant complexes with different paralogs or mutant subunits without the need to suppress production of endogenous proteins (Fig. 1g, h).

We have also developed a nucleosome ejection assay to aid in assessment of CBAF activity and regulation (Fig. 2a, Fig. 4a). The assay is adaptable, easily quantifiable, and requires no specialized equipment. As the force/physics parameters are different, nucleosome sliding activity is not always predictive of ejection activity. Therefore, this assay enables insight into the enzymatic activity of variant complexes that is unattainable with conventional sliding assays (Fig. 2c). In addition, the identity of the nucleosome positioning sequence may be altered to enable investigation of the effects of DNA sequence on nucleosome ejection activity, as has been performed for nucleosome sliding^43^.

This assay provided insight into whether direct ejection or spooling is the predominant mechanism of nucleosome ejection, and if ejection can occur efficiently without histone chaperones. SWI/SNF-family remodelers can transfer histone octamers^44^, exchange H2A/H2B dimers^45^, and disassemble nucleosomes^39^, and were initially thought to require acceptor DNA or chaperones. Disassembly of mononucleosomes has recently been observed in the absence of chaperones or acceptor DNA^27^. Likewise, nucleosome ejection via the spooling mechanism can occur with nucleosome dimer templates in the absence of chaperones^42^. However, the efficiency of direct ejection compared to spooling has not been determined previously in a closed polynuclesome array format, which better resembles *in vivo* chromatin. Ejection from circular polynucleosome arrays without chaperones has also been reported, but the assay cannot distinguish between the two proposed mechanisms^12,27,37^. Here, the ejection assay with T5 exonuclease treatment revealed the clear loss of the final nucleosome from a closed circular array (Fig. 4a). Notably, the assay was performed in the absence of histone chaperones or receptor DNA. Importantly, there is no evidence in the data for a buildup or delay in removal of the final octamer, indicating that direct ejection of the last octamer occurs with similar efficiency as removal of all other octamers on the array. Therefore, while we cannot exclude the possibility that ejection can occur by both direct and spooling mechanisms, the high rate of removal of the final nucleosome indicates that direct ejection is an efficient mode of nucleosome ejection by BAF. Here, interesting future work may be directed at understanding nucleosome modifications or variants that might stimulate or prevent ejection by either of these modes.

Our data confirm that β-actin and BAF53A form an obligate heterodimer within BAF and identify BCL7A as a functional member of the ARP module (Fig. 2c). BCL7A frequently undergoes biallelic inactivation in diffuse large B-Cell lymphoma, and our data identify a mechanistic consequence of its loss in this disease^46^. The results of our investigation into ARPs function align with previous work on a yeast remodeler showing that the ARP module enhances the efficiency with which ATP hydrolysis is coupled to DNA translocation^27^, highlighting the evolutionary conservation of regulatory logic within SWI/SNF family remodelers. This provides additional support for the ability of results obtained using remodelers from different species to predict the roles of homologous subunits and domains in their human counterparts.

The role of ARID1A in CBAF stability and enzymatic activity has been unclear, due to conflicting reports in the literature^4,15,16,31,32^. Here, we show that CBAF complexes readily assemble in the absence of ARID1A and retain WT levels of enzymatic activity, providing compelling evidence for the dispensability of this subunit under our experimental conditions. However, our data do not preclude the possibilities that ARID1A may enhance CBAF assembly efficiency under certain conditions, or that ARID1A may promote chromatin remodeling of variant or modified nucleosomes.

One result of particular interest is the multifaceted regulation of nucleosome ejection by BAF47 through its two stimulatory domains (WH and CTT) and a central autoinhibitory domain (Fig. 4a). Previous studies identified the CTT as a stimulatory domain that potentiates nucleosome sliding and ejection through its interaction with the nucleosome acidic patch^13,35^. In addition, RSC displays enhanced nucleosome ejection activity on nucleosomes with an extended acidic patch^38^. One attractive explanation involves the CTT providing an anchoring point for the complex on the histone octamer (specifically, the ‘dish face’ H2A-H2B dimer) that can assist in chromatin remodeling. However, CBAF complexes lacking BAF47 have only a modest decrease in nucleosome ejection activity—and display an increase in sliding. Therefore, we speculate that the CTT may instead (or additionally) function as a sensor, and stimulate ejection activity when in contact with the nucleosome acidic patch. Notably, the WH domain interacts with the HSA domain of BRG1^30^, which provides the assembly platform for the ARPs^47^, suggesting a possible mechanistic link between BAF47 and the ARP module. Additional studies will be required to determine whether the WH and CTT are purely stimulatory domains, or whether one or both domains instead function to relieve autoinhibition imposed by the central portion of BAF47 (Fig. 4c). Intriguingly, the central portion of BAF47 interacts with BCL7A^30^, which suggests a possible mechanism for BAF47 autoinhibition through modulation of BCL7A, which interacts with the ARP module (Fig 2C).

These data underscore the importance of context when assaying mechanistic contributions. For example, both the ΔARP (Fig. 2h) and the BAF47 ΔWH ΔCTT (Fig. 3f) complexes had decreased ejection activity, but for different reasons. Deletion of the ARP module lowered ejection activity by reducing DNA translocation efficiency, whereas BAF47 truncations decreased ejection via autoinhibition without affecting the core enzymatic activity of BRG1. This raises the possibility that ARP loss may confer an ejection defect on all nucleosomes due to an inherent reduction in motor activity, whereas BAF47 alterations may cause remodeling deficits only on particular variant or modified nucleosomes.

Taken together, our work establishes a versatile and effective system for producing full recombinant CBAF, as well as a novel assay for nucleosome ejection, providing a potent combination for the investigation of CBAF assembly, activity, and regulation. Our results provide insight into subunit assembly dependencies, the regulatory logic governing enzymatic activity, and the fundamental mechanism of nucleosome ejection. For the field, this system can easily be extended to examine mutations linked to cancer and developmental disorders, and adapted for the production of other SWI/SNF(BAF)-family complexes such as GBAF and PBAF—as well as tailored derivatives—for both biochemical and structural studies.

## ACKNOWLEDGEMENTS

This work was supported by U54CA231652 (K.B.J. and B.R.C.) and F30CA225163 (T.S.M.). We thank Tim Formosa, Mahesh Chandrasekharan, Alisha Schlichter, and Margaret Kasten for helpful comments on the manuscript.

## AUTHOR CONTRIBUTIONS

T.S.M. and B.R.C. conceived the study. T.S.M., K.B.J., and B.R.C. designed the experiments. T.S.M, M.L.N., and N.V. performed the experiments. T.S.M. and B.R.C. wrote the manuscript.

## METHODS

### biGBac cloning

The pLibMam vector (pFastBac1-CMV) was a gift from Dr. Erhu Cao. cDNAs encoding each of the CBAF subunits were PCR-amplified using Phusion DNA polymerase in HiFi Phusion buffer (NEB) and subcloned into pLibMam vectors using NEBuilder HiFi (NEB) and homemade chemically competent Top10 *E. coli*. pBig1 and pBig2 expression vectors were assembled using Phusion polymerase and NEBuilder HiFi according to published protocols^24^. DH10B electrocompetent cells (Thermo Fisher) were used for pBig2 cloning.

### Expression vector design

CBAF complexes were assembled using the following plasmids for transfection: pBIG2-WT and pLibMam-ARID1A (WT); pBIG2-WT (ΔARID1A); pBIG2-ΔBCL7A and pLibMam-ARID1A (ΔBCL7A); pBIG2-ΔARP and pLibMam-ARID1A (ΔARP); pBIG2-ΔTetR, pLibMam-BCL7A, pLibMam-BAF53A, pLibMam-DPF2, and pcDNA6-ARID1A (ARID1A-Pulldown). For the DPF2 and BAF47 studies, base expression vectors were pBIG2-ΔBAF47 and pLibMam-ARID1A. Additional pLibMam plasmids encoding DPF2, BAF47, BAF47 ΔWH, BAF47 ΔCTT, and/or BAF47 ΔWH ΔCTT were co-transfected as indicated. A 3xFLAG epitope tag is included on the N-terminus of BAF60A to facilitate purification. A single-chain version of the tetracycline repressor (scTetR) is included on the N-terminus of BAF57 to enable a tet-tethered DNA translocation assay.

### Expression and purification of recombinant CBAF

Expi293F cells (Thermo Fisher) were grown and transiently transfected according to the manufacturer’s instructions. Cells were harvested 72 hours post-transfection by centrifugation for 5’ at 500 g at 4° C. All subsequent steps were performed at 4° C. Cells were washed in TBS, resuspended in Buffer A (20 mM HEPES pH 8.0, 1.5 mM MgCl_2_, 10 mM KCl, 0.25% NP-40, 0.5 mM DTT, protease inhibitors), and incubated on ice 10’ prior to homogenizing with a Dounce (Wheaton or Sigma). Nuclei were pelleted by centrifugation for 5’ at 5000 rpm, resuspended in Buffer C (20 mM HEPES pH 8.0, 25% glycerol, 1.5 mM MgCl_2_, 420 mM KCl, 0.25% NP-40, 0.2 mM EDTA, 0.5 mM DTT, protease inhibitors), Dounce homogenized, and extracted for 30’ with rotation. Nuclear extracts were clarified by centrifugation for 30’ at 20,000 g and applied to anti-Flag M2 affinity gel (Sigma) and rotated for 45’. Flag resin was pelleted by centrifugation for 1’ at 1500 g, washed 3x with Buffer C and 3x with Sizing buffer (20 mM HEPES pH 8.0, 200 mM NaCl, 10% glycerol, 0.5 mM DTT, protease inhibitors). CBAF was eluted 2×30’ with 250 ng/μl 3xFlag peptide (Sigma) in Sizing buffer. Elution fractions were pooled, concentrated to 500-1000 ng/μl using spin concentrators with a 100 kDa cutoff (Amicon), and filtered through a 0.22-micron spin-X column (Corning Costar). Purified, concentrated, and filtered complexes were aliquoted and flash-frozen in liquid nitrogen and stored at −80° C.

### Size exclusion chromatography

Purified complexes were analyzed on a Superose 6 Increase 3.2/300 GL (GE) at 0.01 ml/min in Sizing buffer (20 mM HEPES pH 8.0, 200 mM NaCl, 10% glycerol, 0.5 mM DTT) using an AKTA Pure (GE). Curves in the range of 0.75 ml to 1.75 ml were fitted using Prism as the sum of 2 gaussians corresponding to the aggregated fraction (Void) and the monodisperse fraction (CBAF). The area under each gaussian was calculated according to the equation Area=(Amplitude*Standard Deviation)/0.3989 and used to determine the fraction of CBAF that was monodisperse in each sample.

### ARID1A double purification

CBAF with a 6xHis-tagged ARID1A was purified as described above with the following modifications. Expi293F cells were transfected with the following plasmids: pBIG2-ΔTetR, pLibMam-BCL7A, pLibMam-BAF53A, pLibMam-DPF2, and pcDNA6-ARID1A (ARID1A-Pulldown). After Flag elution, the purified complex was applied to Ni-NTA agarose (Qiagen) and rotated 60’ at 4° C. The resin was washed 3x with Sizing buffer and eluted with 200 mM imidazole in Sizing buffer. The purified complex was concentrated, filtered, and frozen as above.

### Nucleosome assembly

Recombinant *Drosophila* octamers were expressed in BL21-CodonPlus (DE3)-RIL *E. coli*, purified, and assembled into octamers by salt dialysis, as described^27,48^.

200 bp dsDNA fragments with a central Widom 601^49^ nucleosome positioning sequence were produced by digestion of plasmid pUC12×601 with AvaI, purified using a Prep Cell (BioRad) with 4.5% native polyacrylamide gel at 400 V in 0.5x TBE (45 mM Tris-Borate pH 8.0, 1 mM EDTA) and eluted with TE buffer (10 mM Tris, 1 mM EDTA) as described^27^.

Mononucleosomes were assembled at 4° C using salt dialysis by mixing *Drosophila* octamers with DNA at a 1:1 or 1.2:1 molar ratio with 0.1 mg/ml BSA in RB-high (10 mM Tris-HCl pH 7.4, 2 M KCl, 1 mM EDTA, 1 mM DTT) in a Slide-A-Lyzer mini dialysis unit with a 7,000 Da molecular weight cutoff (Thermo Fisher). Assemblies were dialyzed for 1000 minutes against 500 ml of RB-high, which was replaced at a rate of 2 ml/min with RB-low (10 mM Tris-HCl pH 7.4, 50 mM KCl, 1 mM EDTA, 1 mM DTT) using an Econo-pump (BioRad), as described^27,48^.

Nucleosome arrays were assembled using *Drosophila* octamers and the 5 kb pUC12×601 plasmid, which contains 12 repeats of the Widom 601 nucleosome positioning sequence. Arrays were assembled using the same protocol as the mononucleosome assembly described above, but with 3:1, 6:1, 9:1, or 12:1 molar ratios of octamer to plasmid (1:4, 2:4, 3:4 and 4:4 molar ratios of octamer to Widom 601 nucleosome positioning sequence) and a 400 ml starting volume of RB-high.

### ATPase assay

ATPase activity was measured using a colorimetric assay that detects inorganic phosphate by complexation with molybdate-malachite green. Assays were performed under V_max_ conditions at 30° C and 500 RPM in a thermomixer (Eppendorf) by incubation of 200 fmol CBAF with 100 ng of Bluescript plasmid in 5 μl ATPase buffer (10 mM HEPES pH 7.3, 20 mM KOAc, 5 mM MgCl_2_, 5% glycerol, 0.1 mg/ml BSA, 0.5 mM DTT, 1 mM ATP). After 30’, 80 μl MGAM (3 volumes MG (.045% malachite green in 0.1 N HCl) to 1 volume AM (4.3% ammonium molybdate in 4 N HCl)) was added, followed 1’ later with 10 μl 34% w/v sodium citrate. After a 10’ incubation, OD_650 nm_ was recorded, essentially as described^27^.

### DNA translocation assay

A 3 kb plasmid with a tetracycline operator (TetO) was relaxed with *E. coli* topoisomerase I prior to incubation with CBAF. CBAF is anchored to the TetO sequence via the DNA-binding domain of the tetracycline repressor (TetR), which is present as an N-terminal fusion on BAF57. Translocation along the DNA sugar-phosphate backbone produces positive supercoils ahead of the remodeler and negative supercoils in its wake. Topo I only relaxes negative supercoils, resulting in the accumulation of positive supercoils as translocation occurs. DNA translocation experiments were performed as a timecourse using a 50 μl starting volume containing 1250 fmol CBAF, 1250 ng relaxed plasmid, 1 mM ATP, 6.25 U topoisomerase 1 (NEB), and 1 mg/ml BSA in 1xNEB4. Reactions were incubated at room temperature. 9 μl aliquots were removed at each timepoint and heat-inactivated for 20’ at 65° C. Samples were deproteinated with 1 μl 10% SDS and 1 μl 10 mg/ml proteinase K at 50° C for 60’. Samples were ethanol-precipitated and run on a 1.3% agarose gel for 3 hr at 130 V. Gels were stained for 20’ with 1 μl/ml ethidium bromide (EtBr) and scanned on a Typhoon Trio (GE), essentially as described^27^.

### Nucleosome sliding assay

Nucleosome sliding experiments were performed as a timecourse at a 1:8 CBAF to nucleosome molar ratio using a 51 μl starting volume containing 300 fmol CBAF, 2400 fmol nucleosomes, 0.1 mg/ml BSA, and 1 mM ATP in Sliding buffer (10 mM Tris pH 7.5, 50 mM KCl, 3 mM MgCl_2_, 0.5 mM DTT). Reactions were incubated at room temperature or 30° C. At each timepoint, 8.5 μl was removed and quenched by the addition of 100 ng competitor DNA and EDTA to a concentration of 13 mM. Glycerol was added to a final concentration of 10% and samples were loaded on a 4.5% native polyacrylamide gel in 0.4x TBE and run for 50’ at 110 V. Gels were stained for 10’ with 1 μl/ml EtBr and scanned on a Typhoon Trio (GE), essentially as described^27^.

### Nucleosome ejection assay

Nucleosome ejection experiments were performed as a timecourse at a 1:2 CBAF to nucleosome molar ratio (6:1 CBAF:Array molar ratio) in a 120 μl starting volume containing 2400 fmol CBAF, 400 fmol arrays, 6 U *E. coli* Topoisomerase 1 (NEB), 0.1 mg/ml BSA, and 1.25 mM ATP in Sliding buffer (10 mM Tris pH 7.5, 50 mM KCl, 3 mM MgCl_2_, 0.5 mM DTT). Reactions were incubated at 30° C and 500 RPM in a thermomixer (Eppendorf). 20 μl aliquots were removed at each timepoint and heat-inactivated for 20’ at 65° C. Samples were deproteinated with 2.5 μl 10% SDS and 2.5 μl 10 mg/ml proteinase K at 50° C for 60’. Samples were ethanol-precipitated and run on a 0.9% agarose gel for 3 hr at 250 V. Gels were stained for 20’ with 1 μl/ml EtBr and scanned on a Typhoon Trio (GE).

For the ejection assay shown in Fig. 4, the experiment was performed as above with the following modifications: after ethanol precipitation, samples were resuspended in 10 μl NEB4, 10 U T5 exonuclease (NEB) were added, and samples were incubated at 37° C for 60’. Samples were subsequently subjected to gel electrophoresis as above.

### Statistical rigor

Each biochemical assay was performed four times: generally, twice with each of two separate purifications of each variant complex. Within each experiment, 3 replicates (ATPase) or 3-5 linear range timepoints (Translocation, Sliding, Ejection) were used to compare relative activity. Data are normalized to mean WT activity within an assay (ATPase) or to WT activity at each timepoint within an assay (Translocation, Sliding, Ejection). Significance was calculated using a t-test (ATPase) or paired t-test (Translocation, Sliding, Ejection) of each variant complex relative to WT.

