## Supplemental Figures and Legends for "A system for CBAF reconstitution reveals roles for BAF47 domains and BCL7 in nucleosome ejection"

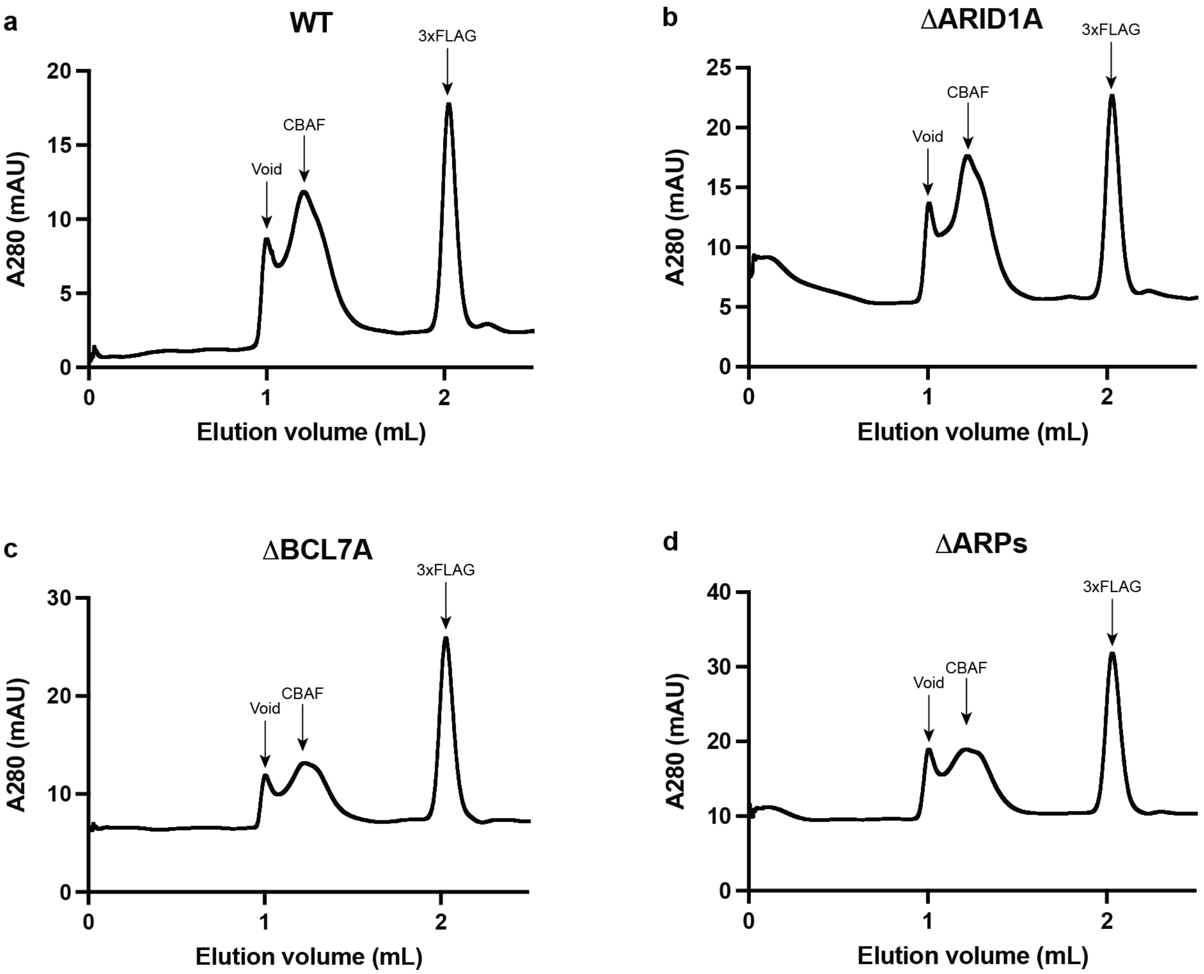


716 **Supplementary Fig. 1, related to Fig. 1.**
**Size exclusion chromatograms for ΔARID1A, ΔBCL7A, and ΔARPs.**
Recombinant CBAF complexes were purified as described and resolved on a Superose 6 increase 3.2/300 at 0.01 mL/min in Sizing buffer.

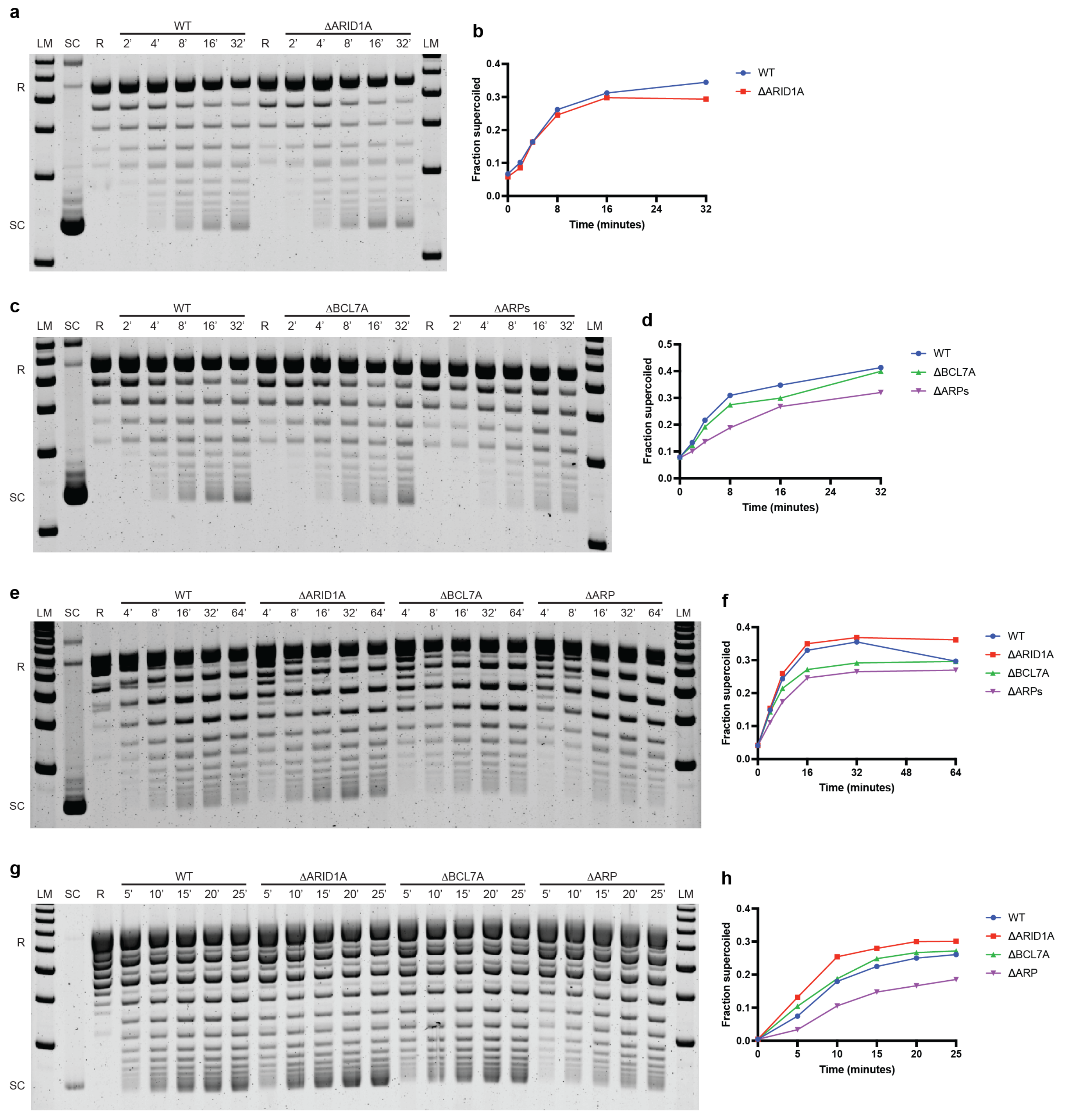


**Supplementary Fig. 2, related to Fig. 2.**
**Additional translocation assay gels and quantifications for ΔARID1A, ΔBCL7A, and ΔARPs.**
Four translocation experiments (**a**, **c**, **e**, **g**) and associated quantifications (**b**, **d**, **f**, **h**). A plasmid with a TetO tetracycline operator was relaxed with *E. coli* topoisomerase I prior to incubation with CBAF variants for the indicated times. CBAF is anchored to the TetO sequence via the DNA-binding domain of TetR. Translocation along the DNA sugar-phosphate backbone produces positive supercoils ahead of the remodeler and negative supercoils in its wake. Topo I only relaxes the negative supercoils, resulting in the accumulation of positive supercoils as translocation occurs, which can be visualized on an agarose
717 gel. R: Relaxed plasmid; SC: Supercoiled plasmid.

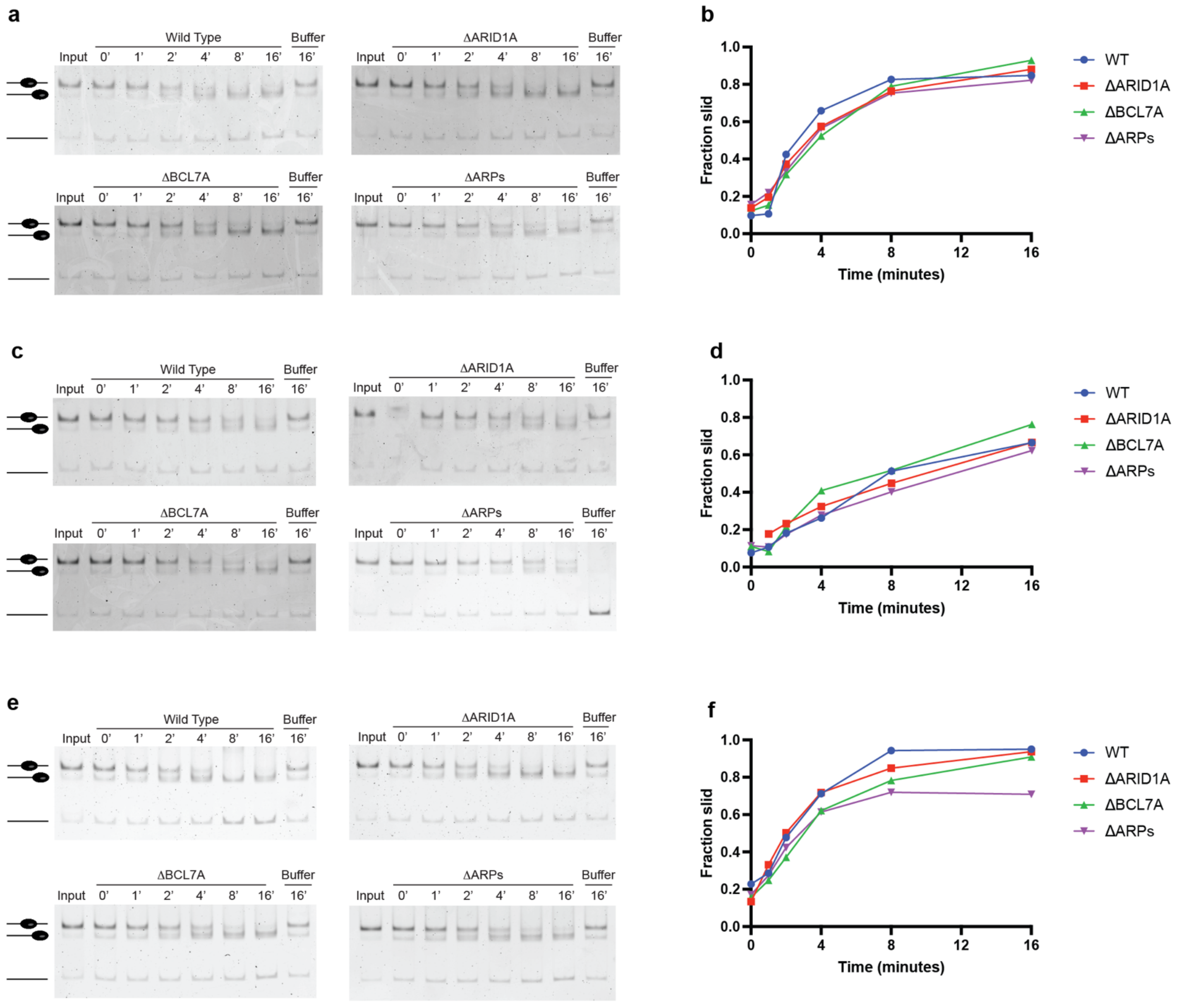


**Supplementary Fig. 3, related to Fig. 2.**
**Additional sliding assay gels for ΔARID1A, ΔBCL7A, and ΔARPs.**
Three sliding experiments (**a**, **c**, **e**) and associated quantifications (**b**, **d**, **f**). Assays performed on a 200 bp Widom 601 nucleosome positioning sequence with *Drosophila* octamers at room temperature (**c**) or 30° C (**a**, **e**).

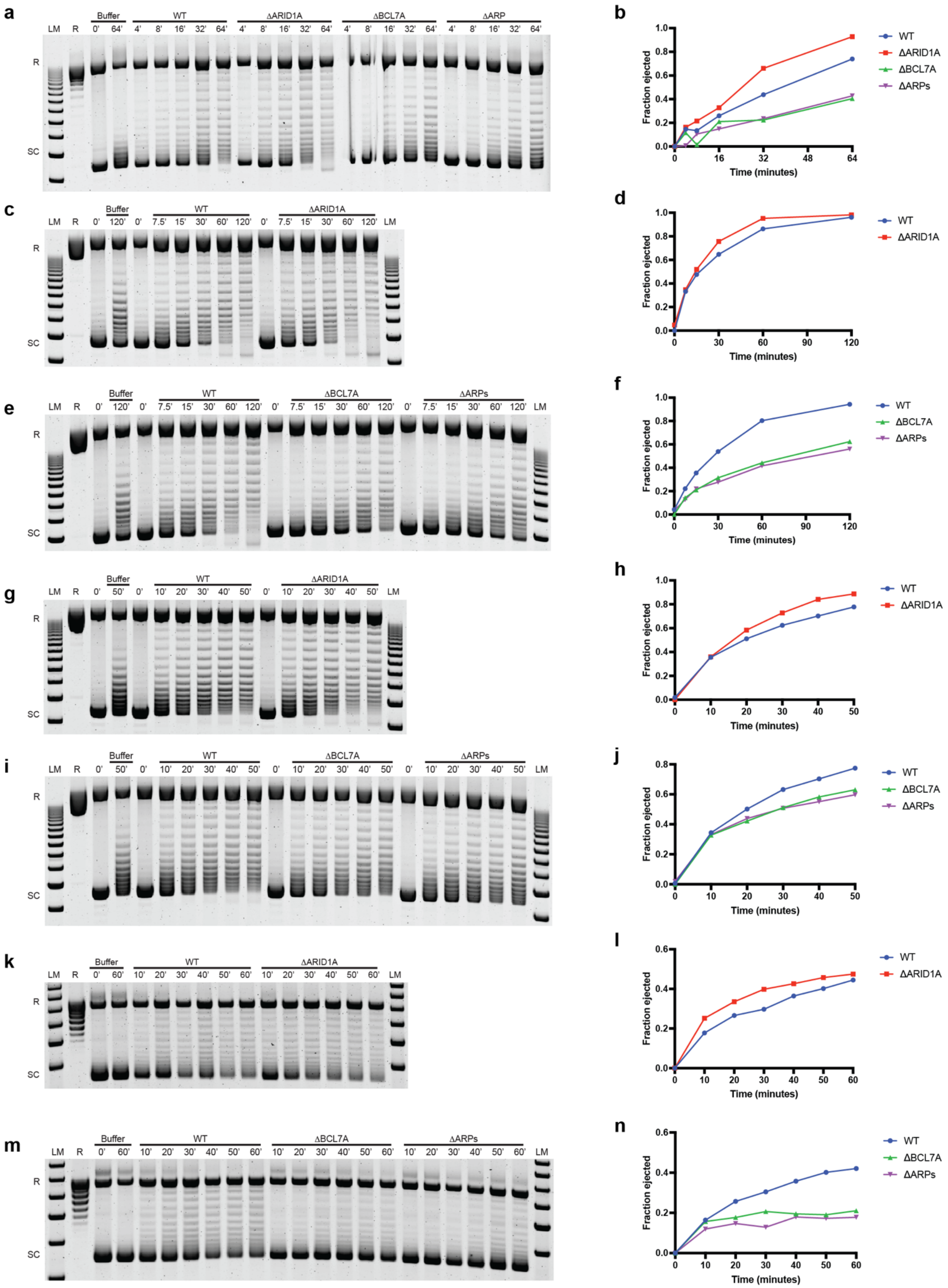


**Supplementary Fig. 4, related to Fig. 2.**
**Additional ejection assay gels for ΔARID1A, ΔBCL7A, and ΔARPs.**
Seven ejection assay experiments (**a**, **c**, **e**, **g**, **i**, **k**, **m**) and associated quantifications (**b**, **d**, **f**, **h**, **j**, **l**, **n**). Arrays were assembled with sufficient histone octamers to saturate all 12 available nucleosome positioning sequences per plasmid, prior to treatment with Topo I. Assembled arrays were incubated with CBAF or Buffer (mock) for the indicated times. R: Relaxed plasmid; SC: Supercoiled plasmid.

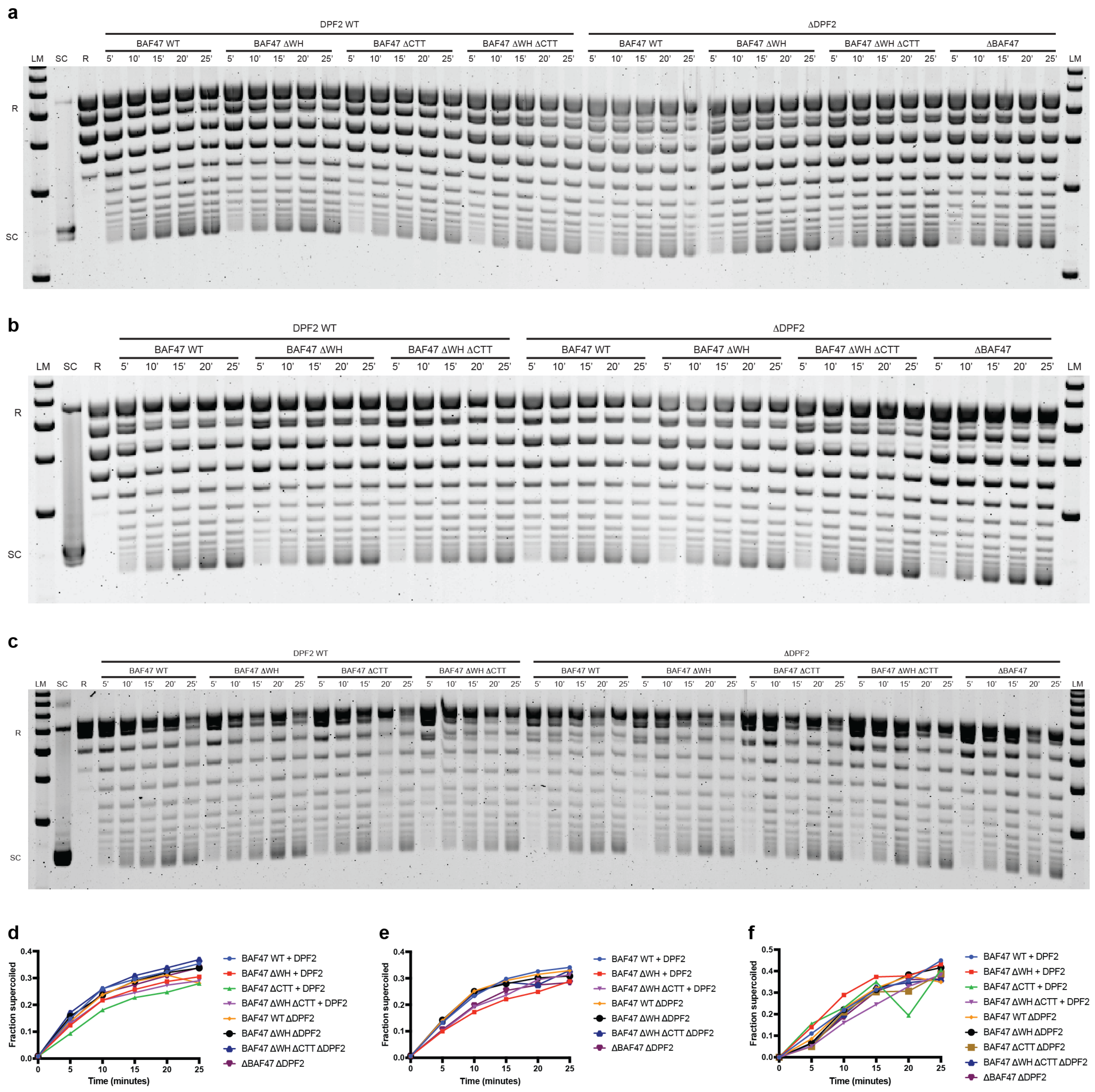


**Supplementary Fig. 5, related to Fig. 3.**
**Additional translocation assay gels for BAF47 and DPF2 truncations and deletions.**
Three additional translocation assays (**a**, **b**, **c**) and quantifications (**d**, **e**, **f**). A plasmid with a TetO tetracycline operator was relaxed with *E. coli* topoisomerase I prior to incubation with CBAF variants for the indicated times. CBAF is anchored to the TetO sequence via the DNA-binding domain of TetR. Translocation along the DNA sugar-phosphate backbone produces positive supercoils ahead of the remodeler and negative supercoils in its wake. Topo I only relaxes the negative supercoils, resulting in the accumulation of positive supercoils as translocation occurs, which can be visualized on an agarose gel. R: Relaxed plasmid; SC: Supercoiled plasmid.

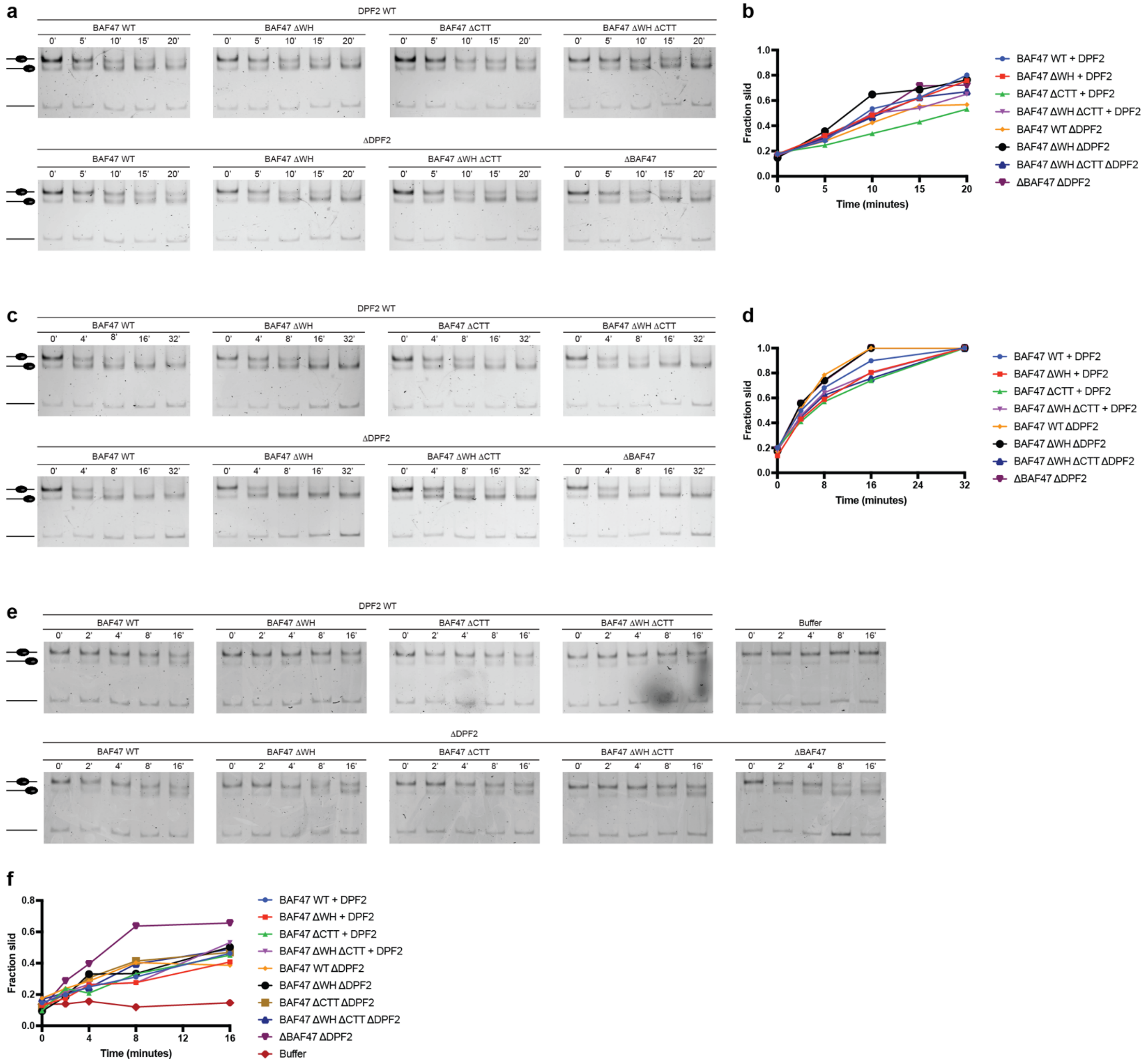


**Supplementary Fig. 6, related to Fig. 3.**
**Additional sliding assay gels for BAF47 and DPF2 truncations and deletions.**
Three sliding experiments (**a**, **c**, **e**) and quantifications (**b**, **d**, **f**). Assays performed on a 200 bp Widom 601 nucleosome positioning sequence with *Drosophila* octamers at room temperature.

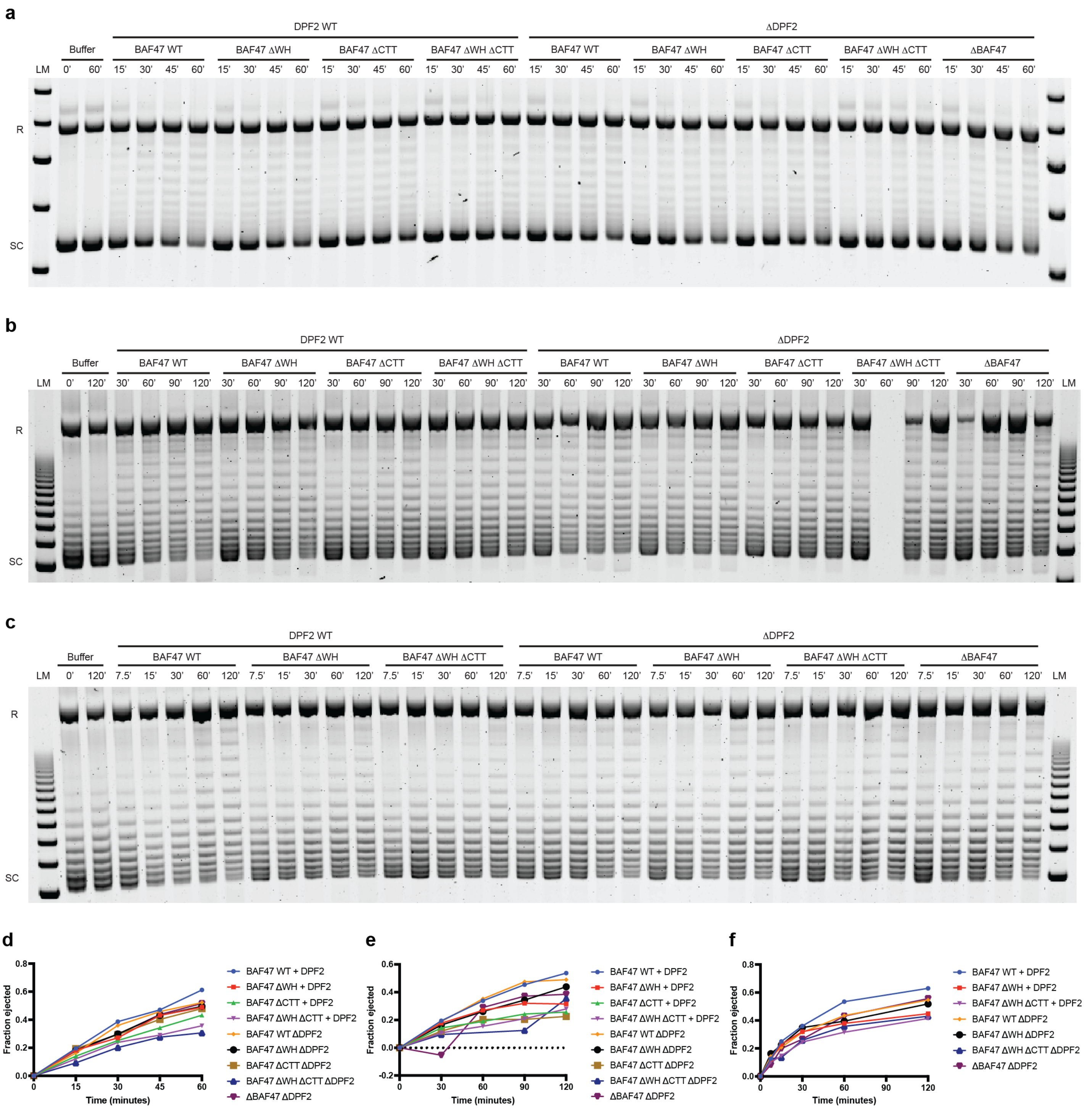


**Supplementary Fig. 7, related to Fig. 3.**
**Additional ejection assay gels for BAF47 and DPF2 truncations and deletions.**
Three ejection assays (**a**, **b**, **c**) and quantifications (**d**, **e**, **f**). Arrays were assembled with sufficient histone octamers to saturate all 12 available nucleosome positioning sequences per plasmid, prior to treatment with Topo I. Assembled arrays were incubated with CBAF or Buffer (mock) for the indicated times. R: Relaxed plasmid; SC: Supercoiled plasmid.
